# Microscopic Motor Alterations in Psychosis and Chronic Cannabis Use

**DOI:** 10.64898/2026.06.25.734496

**Authors:** Filippo Pasqualitto, Alice Tomassini, Angela Muscettola, Cecilia Gabelli, Giovanni Nazzaro, Giovanni Antonio De Bellis, Francesco Torricelli, Gian Marco Gobbi, Maria Giulia Nanni, Luigi Grassi, Luciano Fadiga, Martino Belvederi Murri, Alessandro D’Ausilio

## Abstract

**Background and Hypothesis:** Motor alterations represent an important component of psychotic disorders. Chronic cannabis use, a key risk factor for psychosis, is also associated with sensorimotor dysfunctions. Yet, the hypothesis of a common sensorimotor disturbance remains underinvestigated.

**Study Design.:** In this study, we examined submovements, elementary units of motor output, to search for common subclinical impairments in these populations. Patients with psychosis (n = 17), heavy cannabis users (n = 21), and healthy controls (n = 17) performed a continuous visuomotor synchronization task, consisting in tracking a dot moving on a screen with a finger.

**Study Results.:** Individuals with psychosis and cannabis users exhibited less frequent and more variable submovements compared with healthy controls. Furthermore, when interacting with a pre-recorded human kinematic profile, both groups exhibited attenuated responses to the observed submovements. This alteration was found to be more pronounced in patients with psychosis.

**Conclusions.:** These findings suggest that submovement analysis may reveal subtle, shared alterations in sensorimotor integration in psychosis and chronic cannabis use, providing an objective window onto motor dysfunction not readily captured by current clinical tools.

## 1. Introduction

Recent models of psychotic disorders emphasize that patients may experience impaired separation of self-generated from external sensory data^1,2^. This mechanism is fundamentally based on sensorimotor control^3,4^. Indeed, both early^5^ and more recent descriptions^6–8^ of psychotic disorders suggest that symptoms such as delusions and hallucinations are often accompanied by a variety of neurological soft signs, including subtle motor dysfunctions^9^. Clinical examination include explicit motor signs, generally seen as undesired effects of antipsychotic treatment^10^. Subclinical motor signs, however, may emerge in the earliest stages of the illness among individuals with a high familial risk^11^ or prodromal syndromes^12^ — as well as first-episode patients who have not yet been treated with antipsychotic medication^13,14^. The presence of motor abnormalities independent of antipsychotic treatment suggests that these subtle motor alterations might shed light on the pathophysiology of psychosis^8,9,15^.

Regular high-dose cannabinoid use is also seen in a high proportion of patients with psychotic disorders. Cannabis use may produce similar sensorimotor impairments in the absence of overt psychiatric symptoms^16–22^, while also increasing the risk of developing psychotic symptoms^23–25^, earlier onset of psychosis^23,24,26,27^, as well as various neurobiological and immune changes^28^. Still, it remains unknown how far this similarity extends with respect to basic sensorimotor integration functions in psychosis and chronic cannabis use.

The present study addresses this problem by capitalizing on motor invariants^29^. Motor invariants are recurrent movement features that reflect basic principles of neuromotor control, such as the coordination of pre-programmed (open-loop) and corrective (closed-loop) processes^3,30^. Submovements have been described as discrete, pseudo-rhythmic components (∼2–3 Hz) embedded within continuous movement trajectories^31–33^. They provide insight into the microscopic organization of movement, representing elementary units underlying continuous movement trajectories^34^. From this perspective, complex movements are the result of the successful concatenation of these elementary units, each reflecting the local integration of intrinsic motor commands and extrinsic sensory feedback during motor execution^35–37^. The presence, timing, and regularity of submovements are increasingly recognized as informative biological correlates of sensorimotor integration^38–43^. At present, however, it remains unknown whether they may be found altered in the context of psychotic disorders and chronic cannabis use. The detection of abnormalities of motor invariants in these populations may provide a fine-grained description of the pathophysiology of sensorimotor impairment.

## 2. Methods

### 2.1 Participants

Patients were recruited from the inpatient psychiatric unit of Sant’Anna Hospital (Ferrara, Italy). Three groups were enrolled in parallel: patients with psychotic disorders (PwP), healthy controls (HC), and healthy regular cannabis users (HC-CU).

PwP were required to have experienced an acute psychotic episode within 1 to 10 days prior to data collection and to be receiving antipsychotic medication at the time of testing. All participants were right-handed and had normal or corrected-to-normal vision. HC and HC-CU completed a DSM-5 psychosis screening; only those with no evidence of current or lifetime psychotic disorders were included. HC-CU were additionally required to report regular cannabis use over the preceding three months; individuals with self-reported problematic use or a history of treatment-seeking suggestive of cannabis use disorder were excluded. Participants who failed to meet minimum task compliance criteria were also excluded.

Initial enrolment yielded 19 PwP, 17 HC, and 22 HC-CU (Table 1). Two participants from the PwP group and one from the HC-CU group were subsequently excluded due to poor task compliance, resulting in final sample sizes of n = 17 (PwP; mean age 34.0, SD 13.0), n = 17 (HC; mean age 31.7, SD 9.8), and n = 21 (HC-CU; mean age 25.0, SD 4.1). Full demographic details are reported in Supplementary Materials, Table S.1.

**Table 1.** Demographic characteristics of the participants. Sex counts and mean age in years with standard deviation (SD) are reported for each group. Abbreviations: HC, healthy controls; HC-CU, healthy controls who are regular cannabis users; PwP, patients with psychosis.

| Group | Sex | Mean age (SD) |
| --- | --- | --- |
| HC ($n = 17$) | 11F, 6M | 31.7 (9.8) |
| HC-CU ($n = 21$) | 6F, 15M | 25 (4.1) |
| PwP ($n = 17$) | 10F, 7M | 34 (13) |

All participants provided written informed consent (Approval: EM255-2020-UniFe/170592-EM) following explanation of the task and procedures, in accordance with local ethics committee guidelines and the Declaration of Helsinki.

### 2.2 Behavioral task

Participants completed a visuomotor synchronization task (Figure 1, a–b), performing rhythmic flexion-extension movements of the right index finger at the metacarpophalangeal joint, synchronized with a dot moving on a computer screen. Participants were instructed to synchronize either in-phase (moving in the same direction as the dot) or anti-phase (moving in the opposite direction), with the ulnar side of the forearm resting on a comfortable support. The visual stimulus followed one of two trajectories: a non-biological, perfect sinusoidal trajectory, or a biological, quasi-sinusoidal trajectory based on pre-recorded human kinematic trace (HKT)^38^. The sinusoidal stimulus had an amplitude comparable to the mean amplitude of the biological trajectories (frequency: 0.25 Hz). Each participant completed three conditions in randomized order: sinusoidal in-phase, biological in-phase, and biological anti-phase. Each condition comprised three continuous 30-second blocks (nine blocks total), preceded by a familiarization block. Stimuli were displayed using VLC media player on a secondary monitor.

**Figure 1.**
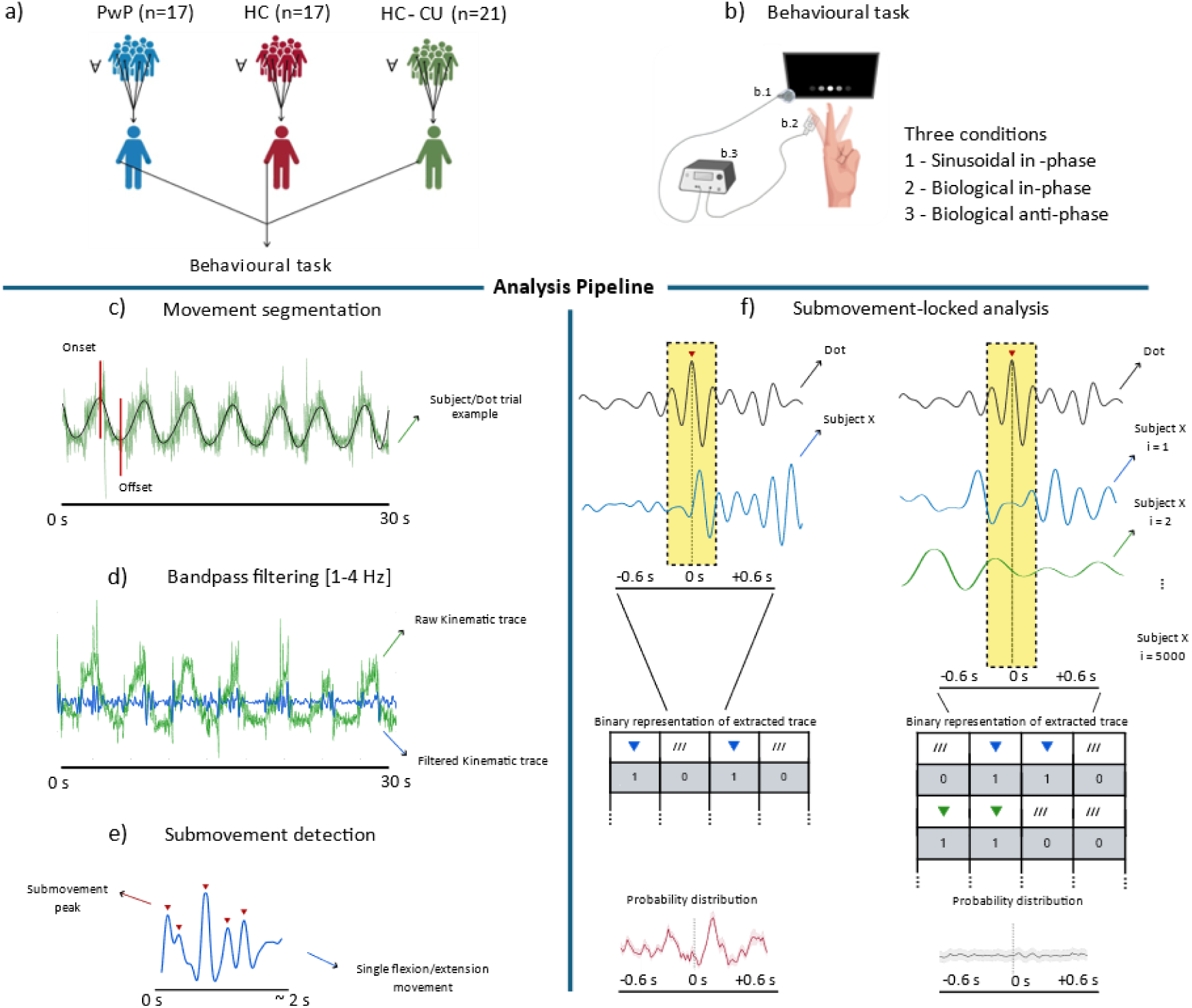
Overview of the experimental design and kinematics’ signal processing. (A) Three groups of participants were tested: individuals with psychosis (PwP, n = 17), healthy controls (HC, n = 17), and healthy cannabis users (HC-CU, n = 21). (B) All participants (∀) performed a visuomotor synchronization task with three conditions: sinusoidal in-phase, biological in-phase, and biological anti-phase. The experimental set-up included an Arduino-based microcontroller (B.3) placed within a resistant container and connected to an accelerometer (B.2; LSM6DS3 sensor). A photodiode (B.1) was positioned at the edge of the monitor to measure the onset and offset of visual stimuli. (C) Timestamps corresponding to the initiation of movements by the subject and the human pre-recorded kinematic trace (HKT) were identified. The figure shows an example of raw kinematic trace recorded over a 30 s block. An example of movement onset and offset is shown using vertical red lines. The green signal represents the raw kinematic data, while the bold line indicates the low-pass filtered signals (cutoff frequency: 0.6 Hz), showing the task-instructed rhythmic movements at 0.25 Hz. This filtered signal was only used for timestamps selection. (D) A fourth-order Butterworth bandpass filter (1–4 Hz) was applied to focus on the frequency range for submovements. The figure shows an example of raw kinematic trace during a 30 s block (green signal) with an example of the same, filtered trace, superimposed to it (blue signal). (E) The figure shows an example of filtered kinematic data relative to a single movement. Submovements were detected (little red arrows pointing downward) by identifying local peaks in the bandpass filtered kinematic signals. (F) The left side of the panel shows time-locked kinematics extraction: the peaks in the HKT trace served as reference points for a time-locked extraction of participants’ kinematics within a window (from -0.6 s to 0.6 s) highlighted by the shaded yellow area. The right side of the panel shows surrogate kinematic extraction where kinematic data were extracted (using the same time-window) by randomly shuffling participants’ movements across 5,000 iterations. For both real and surrogate data, each extracted trace was converted into a binary representation indicating the presence (1) or absence (0) of a submovement at each timepoint. These binary matrices were then used to compute probability distributions of submovement occurrence.

**Figure 2.**
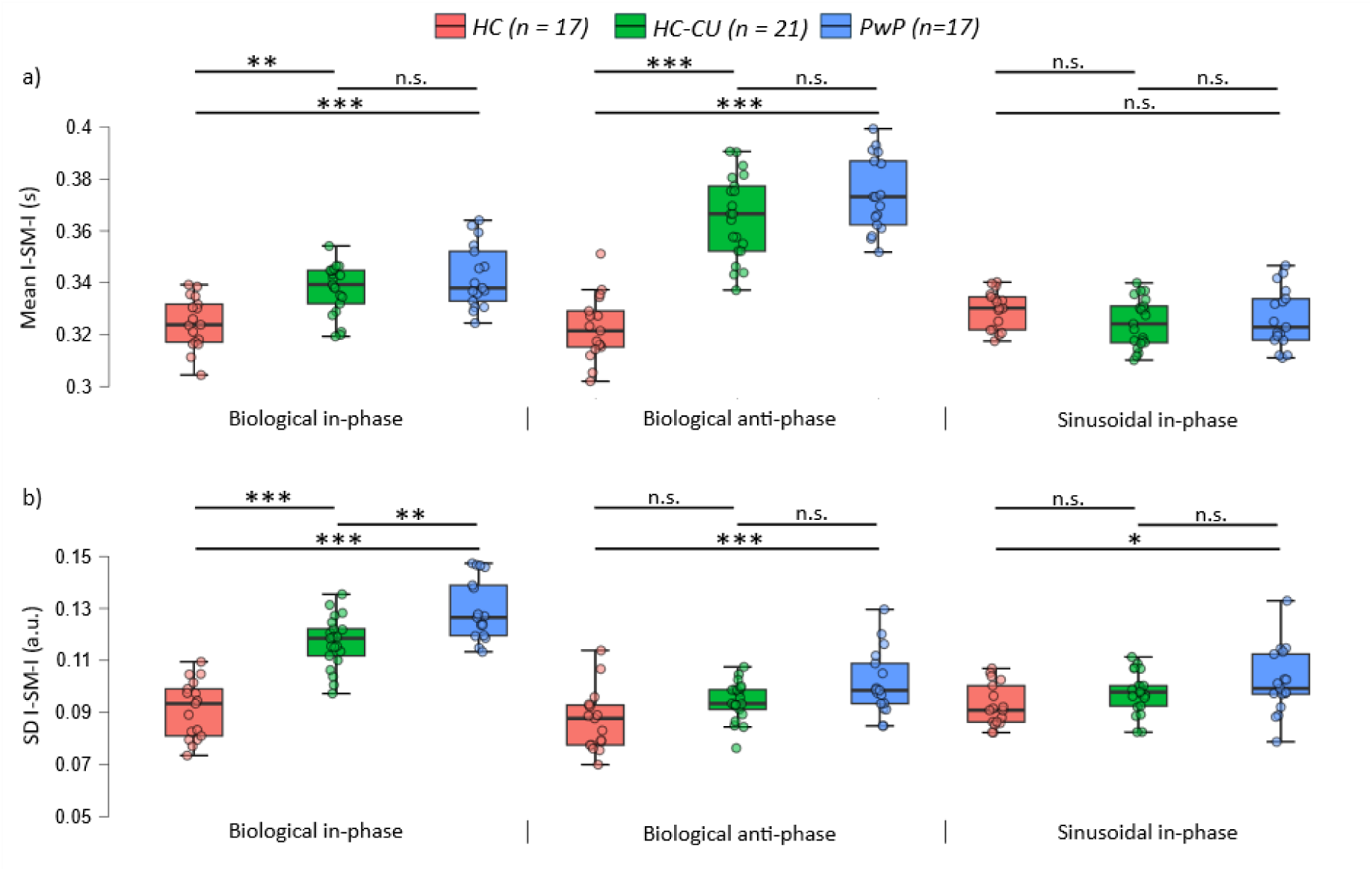
Inter-submovement intervals (I-SM-I) across groups and experimental conditions. Mean values (A) and variability expressed as SD (B), for all groups (HC = Healthy Controls, HC-CU = Healthy Controls – (regular) Cannabis Users, and PwP = Patients with Psychosis) in the three conditions (biological in-phase, biological anti-phase, and sinusoidal in-phase). Significance levels are represented as: n.s. = non-significant, *p < .05, **p < .01, ***p < .001.

### 2.3 Kinematic data recording

Kinematic data were recorded using a custom-built Arduino-based system connected to a 6-degree-of-freedom inertial measurement unit (IMU) integrating a 3-axis accelerometer and a 3-axis gyroscope (LSM6DS3 sensor; MathWorks documentation: <u>link</u>). Signals were sampled at 100 Hz; analyses focused on the z-axis, corresponding to index finger flexion-extension. The sensor was attached to the tip of the right index finger using an adhesive strap. Custom MATLAB code handled stimulus presentation, data acquisition, screen synchronization, and calibration. Setup and sensor placement were kept consistent across all participants and groups.

### 2.4 Kinematic data analysis

All analyses were conducted using custom MATLAB code and the FieldTrip toolbox^44^.

1. Movement segmentation. Continuous recordings were segmented into nine blocks. Raw accelerometer data were downsampled to 60 Hz to match the screen refresh rate and further segmented into individual flexion/extension movements (example of single movement’s onset/offset in Figure 1,c). Basic synchronicity was verified by computing the mean absolute difference between participants’ and HKT timestamps (see Supplementary Material, Section 5).
2. Submovement detection. Raw acceleration signals were bandpass-filtered (1–4 Hz) using a fourth-order, two-pass Butterworth filter (Figure 1,d). Submovements were identified as local peaks in the filtered segments, for both participant data and the HKT trace (Figure 1,e). The mean and standard deviation (SD) of the inter-submovement interval were computed for each condition, pooled across blocks, and averaged within each participant.
3. Submovement-locked analysis. For the biological conditions, participants’ acceleration traces were segmented from −0.6 to 0.6 s around HKT submovements and averaged to examine the temporal relationship between participants’ and HKT submovements (Figure 1,f). The probability of generating a submovement relative to HKT submovements was estimated by counting peaks at each timepoint and dividing by the total number of peaks in the analyzed segments. Probabilities were binned (33 ms, non-overlapping) and expressed as percentage deviations from the mean probability over the full window.

### 2.5 Statistical analysis

1. Submovement periodicity. The mean and SD of inter-submovement intervals were analyzed using separate mixed-design ANOVAs with Group (HC, HC-CU, PwP) as the between-subject factor and Condition (sinusoidal in-phase, biological in-phase, biological anti-phase) as the within-subject factor, implemented in JASP (Version 0.19.3). Bonferroni-corrected post hoc comparisons were applied where relevant.
2. Submovement-locked modulation. To test whether submovement occurrence probability was significantly modulated relative to HKT submovements, observed probabilities were compared against surrogate data (5000 iterations; Figure 1,f) in which the temporal association between participant and HKT movements was disrupted. At each timepoint, a two-tailed paired-sample t-test was applied, with False Discovery Rate (FDR) correction (q = 0.05) using the Benjamini-Hochberg procedure^45^, applied separately per condition and group. Group differences in modulation amplitude and latency were then assessed within the biological in-phase condition which, unlike the biological antiphase condition, showed significant modulation compared with surrogate data (in line with previous findings on biological in-/anti-phase condition^37^). This analysis was performed over the 0–0.6 s post-submovement window using one-way ANOVAs, with the largest peak per participant compared across groups for both submovement-locked acceleration and occurrence probability.
3. Association with clinical and pharmacological variables. In the PwP group, associations between submovement features and clinical or pharmacological variables were examined in three steps, controlling for cannabis use. Submovement features included inter-submovement interval mean and SD across all conditions, and peak amplitude and latency from submovement-locked acceleration and probability in the biological in-phase condition. Clinical profiles were assessed using the Positive and Negative Syndrome Scale (PANSS^46^), the Brief Psychiatric Rating Scale (BPRS^47^), and the St. Hans Rating Scale (SHRS^48^) for extrapyramidal symptoms. Pharmacological profiles via chlorpromazine-equivalent dosage. Linear regressions first tested whether cannabis use (yes/no) predicted clinical or pharmacological measures and submovement features. Zero-order Pearson correlations were then computed between submovement features and clinical or pharmacological variables, followed by partial correlations controlling for cannabis use. FDR correction (Benjamini-Hochberg, q = 0.05) was applied within each family of tests.

## 3. Results

### 3.1 Participant characteristics

Clinical characteristics of PwP were assessed through PANSS, BPRS and SHRS psychometric instruments (Table 2). The mean total PANSS score was 69.1, falling within the range typically associated with mild severity (cut-off scores: 58–74), and the mean BPRS total score was 41.5, consistent with moderate severity (cut-off scores: 40–45). No participant showed clinically significant extrapyramidal symptoms (SHRS cut-off score ≥ 2; 48). Antipsychotic dosages varied substantially across individuals, ranging from 20 to ∼6000 mg/day (Supplementary Materials, Table S.2). No participant in the HC or HC-CU groups reported psychiatric symptomatology at anamnestic assessment. In the HC-CU group, cannabis use disorder was ruled out through both a structured interview and the Cannabis Experience Questionnaire^49^.

**Table 2.**
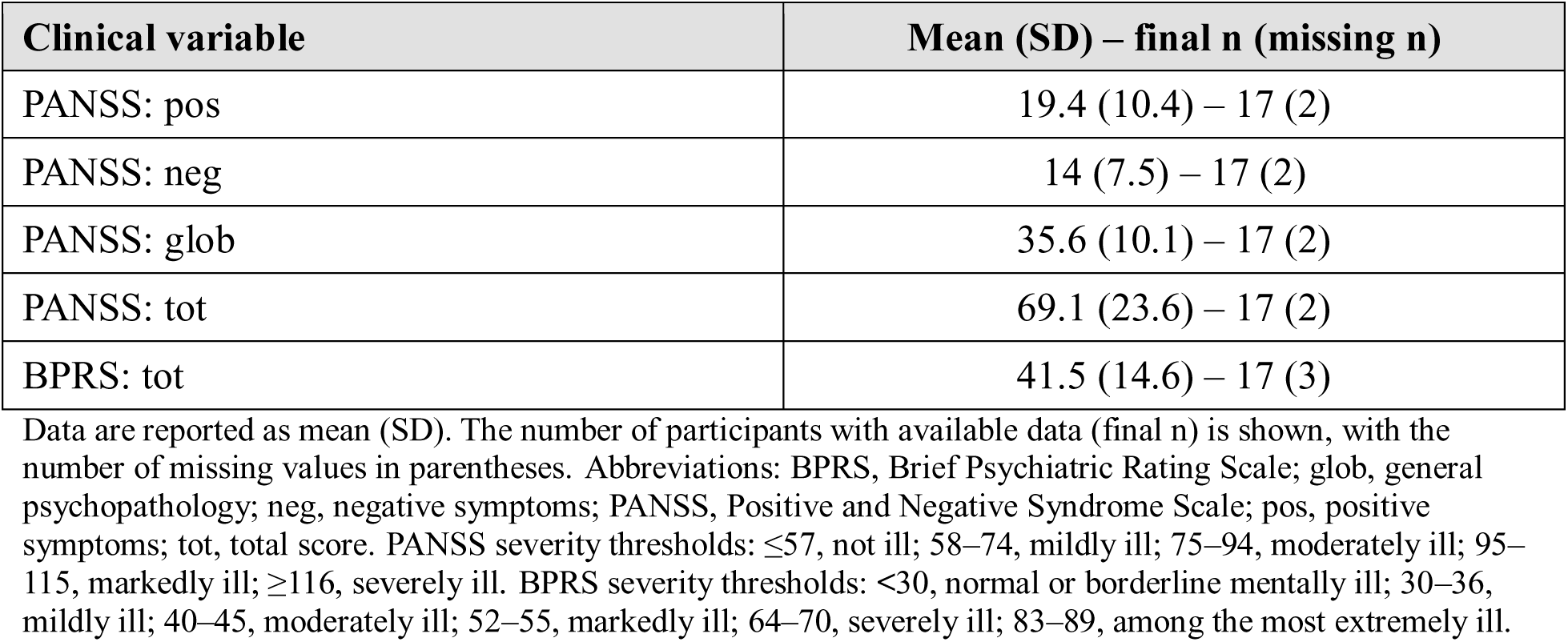
Clinical characteristics of patients with psychosis. Data are reported as mean (SD). The number of participants with available data (final n) is shown, with the number of missing values in parentheses. Abbreviations: BPRS, Brief Psychiatric Rating Scale; glob, general psychopathology; neg, negative symptoms; PANSS, Positive and Negative Syndrome Scale; pos, positive symptoms; tot, total score. PANSS severity thresholds: ≤57, not ill; 58–74, mildly ill; 75–94, moderately ill; 95–115, markedly ill; ≥116, severely ill. BPRS severity thresholds: <30, normal or borderline mentally ill; 30–36, mildly ill; 40–45, moderately ill; 52–55, markedly ill; 64–70, severely ill; 83–89, among the most extremely ill.

### 3.2 Altered submovement periodicity in patients with psychosis and regular cannabis users

The mean inter-submovement interval showed significant main effects of Group (*F*(2, 52) = 23.525, *p* < .001, *η²* = .188) and Condition (*F*(2, 104) = 162.744, *p* < .001, *η²* = .291), and a significant Group × Condition interaction (*F*(4, 104) = 61.343, *p* < .001, *η²* = .220). Both PwP and HC-CU produced submovements at lower frequency (longer intervals) than HC, particularly in the biological conditions. Intervals were shorter in the sinusoidal than in the biological conditions for both PwP (sinusoidal in-phase vs. biological in-phase: *t* = -5.875; *p* <.001; *Cohen’s d* = -1.327; sinusoidal in-phase vs. biological anti-phase: *t* = -17.336; *p* <.001; *Cohen’s d* = -4.049) and the HC-CU group (*t* = -5.33; *p* <.001; *Cohen’s d* = -1.12; *t* = -16.4; *p* <.001; *Cohen’s d* = -3.446). In contrast, the HC group showed no difference between the sinusoidal in-phase and biological in-phase condition (*p* = .232) and even displayed the opposite pattern, with significantly longer intervals in the sinusoidal in-phase than in the biological anti-phase condition (*t* = 2.618; *p* = .031; *Cohen’s d* = .611). Within-condition group comparisons confirmed that both PwP and HC-CU had significantly longer inter-submovement intervals – indicating a lower emission frequency – than HC in both biological conditions (in-phase: HC vs PwP, *t* = -4.366, *p* < .001, *Cohen’s d* = -1.497; HC vs. HC-CU, *t* = -3.354, *p* = .003, *Cohen’s d* = -1.094; anti-phase: HC vs PwP, *t* = -12.737, *p* < .001, *Cohen’s d* = -4.369; HC vs. HC-CU, *t* = -11.081, *p* < .001, *Cohen’s d* = -3.615), with no significant differences between PwP and HC-CU (in-phase: *p* = .658; anti-phase: *p* = .069). All groups performed comparably in the sinusoidal condition (all *p* ≥ .534).

Inter-submovement interval variability showed significant effects of Group (*F*(2, 52) = 27.867, *p* < .001, *η²* = .259), Condition (*F*(2, 104) = 81.772, *p* < .001, *η²* = .241), and their interaction (*F*(4, 104) = 18.040, *p* < .001, *η²* = .106). The PwP group showed significantly greater variability than HC across all conditions (sinusoidal in-phase: *t* = 2.538, *p* = .037, *Cohen’s d* = .87; biological in-phase: *t* = 10.784, *p* < .001, *Cohen’s d* = −3.699; biological anti-phase: *t* = 4.019, *p* < .001, *Cohen’s d* = 1.378). The HC-CU group showed intermediate variability, differing significantly from both HC and PwP only in the biological in-phase condition (HC vs. HC-CU: *t* = −7.652, *p* < .001, *Cohen’s d* = −2.497; PwP vs. HC-CU: *t* = 3.695, *p* = .001, *Cohen’s d* = 1.202), with no differences in the remaining conditions (all *p* ≥ .092). Variability remained stable across conditions in HC (all *p* ≥ .116), whereas both PwP and HC-CU showed a marked increase in variability in the biological in-phase condition relative to both the biological anti-phase (PwP: *t* = 10.317, *p* < .001, *Cohen’s d* = 2.699; HC-CU: *t* = 9.180, *p* < .001, *Cohen’s d* = 2.161) and sinusoidal in-phase conditions (PwP: *t* = 10.166, *p* < .001, *Cohen’s d* = 2.659; HC-CU: *t* = 7.912, *p* < .001, *Cohen’s d* = 1.862).

Altogether, these findings indicate that PwP and HC-CU are associated with alterations in both submovement periodicity and timing variability, which are most evident when tracking biological motion.

### 3.3 Patients with psychosis and regular cannabis users exhibit reduced alignment with observed submovements

During in-phase synchronization, participants’ kinematics exhibited a prominent peak approximately 180 ms after the dot’s submovements, with smaller peaks at roughly ±400 ms reflecting intrinsic submovement periodicity (Figure 3,a); this modulation was markedly reduced during anti-phase synchronization (Figure S.1,a). The pattern was qualitatively consistent across groups and with previous findings^38^. Submovement occurrence probability confirmed a statistically significant deviation from surrogate data at approximately +180 ms in all groups during in-phase (Figure 3,b), but not anti-phase, synchronization (Figure S.1,b). We therefore focused on the in-phase condition to examine whether the observed modulation differed across groups in either amplitude or latency.

**Figure 3.**
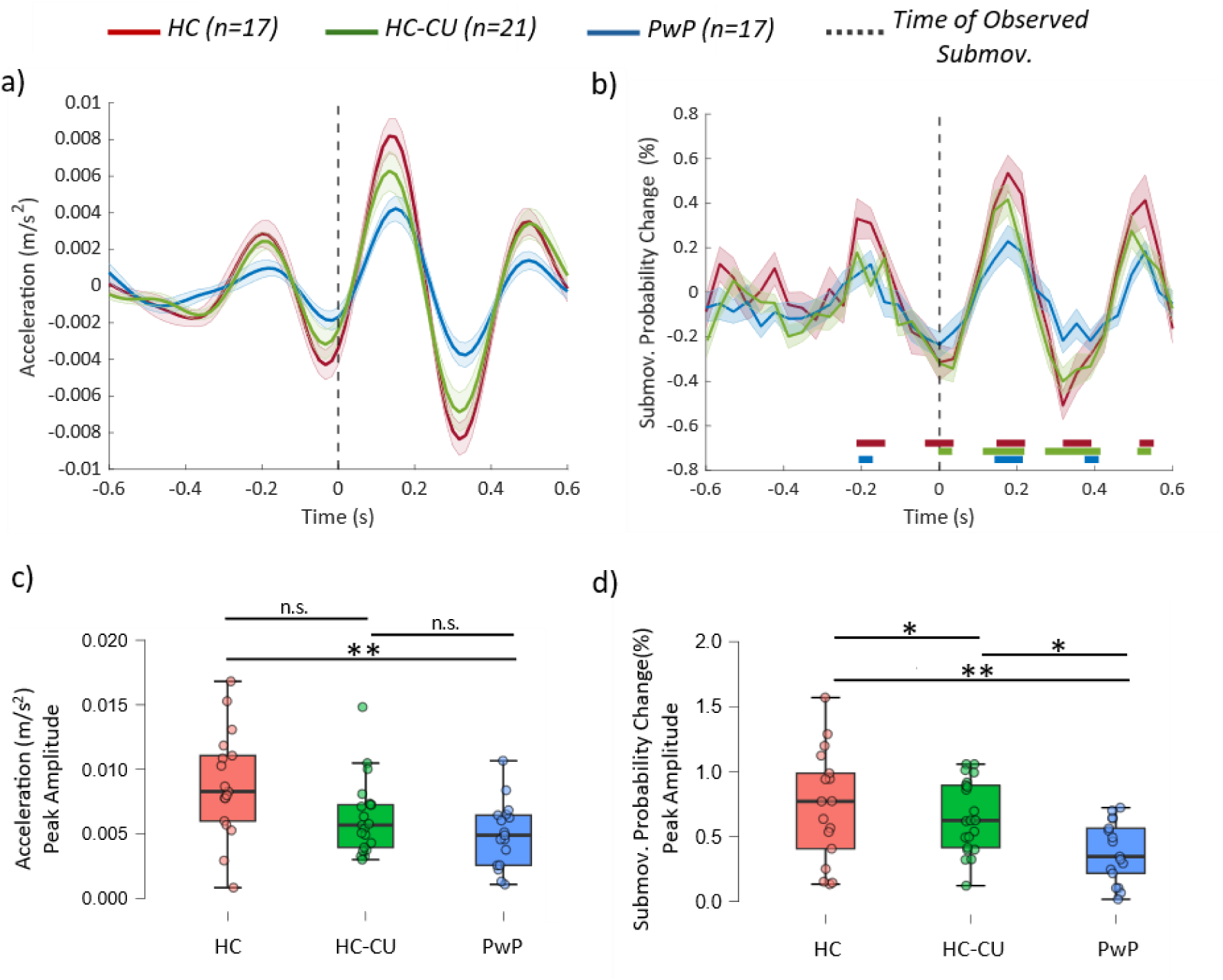
Submovement-locked kinematics and probability across groups in the biological in-phase condition. Acceleration profiles (A) and, probability of submovement occurrence (B), expressed as percentage deviation from the mean over the ±0.6 s interval. In probability time-series, horizontal bars indicate intervals where the original data significantly differ from the surrogate distribution (after FDR correction). Boxplots illustrate peak amplitude modulation for the acceleration profiles (C) and the submovement probability (D) across all groups. Significance levels are represented as: n.s. = non-significant, *p < .05, **p < .01, ***p < .001. Group abbreviations: HC = Healthy Controls, HC-CU = Healthy Controls – (regular) Cannabis Users, and PwP = Patients with Psychosis.

Group comparisons in the in-phase condition revealed no significant kinematic differences in modulation latency (estimated via subject-wise peak detection between 0 and 0.6 s; see Methods; *F*(2, 52) = .723, *p* = .490, *η²* = .027), but a significant group effect on amplitude (*F*(2, 52) = 6.751, *p* = .002, *η²* = .206; Figure 3,c). Amplitude was largest in HC and significantly greater than in PwP (*t* = 3.633, *p* = .002, *Cohen’s d* = 1.246). The HC-CU group showed intermediate amplitude, which did not differ significantly from either the HC (*p* = .063) or PwP group (*p* = .468). The probability of producing a submovement following the dot’s submovement also differed significantly across groups (*F*(2, 52) = 13.833, *p* < .001, *η²* = .347; Figure 3,d), with PwP showing the lowest and HC-CU intermediate values; all pairwise comparisons were significant (HC vs. PwP: *t* = 5.26, *p* < .001, *Cohen’s d* = 1.804; HC vs. HC-CU: *t* = 2.774, *p* = .023, *Cohen’s d* = .905; HC-CU vs. PwP: *t* = 2.756, *p* = .024, *Cohen’s d* = .899). No significant group differences in modulation latency were observed (*F*(2, 52) = 2.983, *p* = .069, *η²* = .103).

Altogether, when exploring submovement alignment to external sensory events, patients with psychosis demonstrated an attenuation evident in both kinematic and probability indices, whereas cannabis users exhibited a milder reduction restricted solely to response probability relative to healthy controls.

### 3.4 No association between submovement features and clinical/pharmacological variables

Among PwP, cannabis use did not significantly predict any clinical/pharmacological variable (all *FDR-corrected p* ≥ .820; Supplementary Materials, Table S.6) or submovement features in any condition (all *FDR-corrected p* ≥ .078; Table S.7). Zero-order Pearson correlations revealed a tendency for longer inter-submovement intervals to be associated with greater symptom severity — particularly for positive symptoms and global psychopathology — but none survived FDR correction (all *FDR-corrected p* ≥ .310; Table S.8). Partial correlations controlling for cannabis use yielded the same pattern (all *FDR-corrected p* ≥ .356; Table S.9).

## 4. Discussion

This study examined the properties of submovements — elementary units of motor output — as a novel, robust assessment of sensorimotor control^35,38,43^. We found that submovement emission frequency was higher and timing more regular in healthy participants compared with chronic cannabis users and individuals with psychosis. Individuals with psychosis showed the greatest temporal variability, differing significantly from both healthy controls and cannabis users. This finding supports the notion of a dysregulation in mechanisms underlying sensorimotor integration. Additionally, when quantifying the ability to micro-align motor outputs to external sensory events (i.e., the dot’s motion), patients with psychosis and, to a lesser extent, regular cannabis users performed consistently worse than healthy controls. Taken together, these results reveal alterations in an otherwise robust feature of movement organization^35,36,38–40,50,51^. The pattern of findings suggests greater similarity between patients with psychosis and cannabis users than between cannabis users and healthy controls. This impairment may reflect shared genetic or environmental vulnerabilities that disrupt sensorimotor integration, potentially contributing to the development of psychosis. At the same time, submovement features in psychosis were not associated with pharmacological treatment or clinical scales. This dissociation suggests that submovement alterations capture aspects of sensorimotor dysfunction not reflected in standard clinical assessments, potentially offering a complementary and objective measure of a basic neurobiological deficit.

In fact, much remains to be characterized beyond overt clinical signs, particularly in the domain of subclinical motor alteration^9,52–54^. These subtle abnormalities may include disrupted temporal coordination^55^, impaired adaptation to contextual changes^56^, and reduced integration of sensory information when predicting the consequences of others’ actions^57^. However, the tasks used in those studies engage overlapping cognitive and sensorimotor processes.

In contrast, the organization of submovements represents a low-level, intrinsic property of motor control^34,35^ with temporal characteristics that reflect fundamental sensorimotor integration processes^36,58^. Our data, extending beyond previous studies suggesting subclinical motor, pre-cognitive disfluencies in individuals with psychosis^59,60^, suggest that these patients may rely disproportionately on open-loop motor strategies driven primarily by internal motor commands, with diminished sensory-based adjustments. This may represent a micro-analytic manifestation of impaired integration of sensory evidence into prediction errors, which could lead to an insufficient or unreliable updating of prior motor plans in response to accumulating sensory information^1,2,61–63^. Functionally, this manifests as reduced flexibility in the reorganization of motor outputs, which may impair the ability to adapt to dynamic and unpredictable environments^1,2,61^.

While the action of cannabinoids have been investigated for a potential therapeutic effect in certain movement disorders — i.e., Parkinson’s disease or Huntington’s disease^64^ — the chronic use of cannabis has been associated with sensorimotor impairments during cognitive-motor tasks^21^ and with alterations in brain networks involved in motor control^16,17,65^. In the present study, the cannabis users group showed a milder pattern of submovement-level dysregulation compared with individuals with psychosis, laying between the healthy control and patient groups. This is consistent with the observation that motor alterations in chronic cannabis users, while documented, are variable, not always replicated^66^, and likely less pronounced than those seen in psychosis. It is also important to note that the risk of developing schizophrenia or other psychotic disorders is up to 8.9% in individuals who use cannabis, compared with 0.6% in non-users^67^, suggesting that cannabis use may contribute to triggering psychosis, but only in a subset of individuals. Future research with larger cohorts will be needed to determine whether subgroups of cannabis users can be distinguished based on their submovement profiles.

These findings suggest that individuals with psychosis and cannabis users may rely disproportionately on pre-programmed, open-loop strategies, even in tasks requiring continuous, feedback-based adaptation. The core impairment may consist in reduced weighting of sensory feedback, potentially reflecting learned “distrust” in its reliability due to increased uncertainty/imprecision^68^. This may be associated with a more general alteration in the calibration between accuracy and confidence, as supported by evidence that patients with psychosis maintain high confidence despite reduced performance during early integration phases^57^. In line with the “jumping to conclusions” bias commonly reported in psychosis^69^, such miscalibration may reinforce a tendency to form unstable or premature beliefs. These impairments may parallel broader inferential dysfunctions in psychosis, in which distorted beliefs persist despite conflicting sensory input, reflecting an imbalance in precision weighting between prior beliefs and incoming sensory evidence^61–63^.

The current findings may inform efforts to identify objective motor-based indices of psychosis and at-risk states^70^. Accelerometer-based tools have been used to quantify gross motor activity and circadian rhythms in clinical populations^71^, while fine-grained analyses of submovement-level dynamics remain rare^72^, to our knowledge. Our results indicate that alterations in the microstructure of motor output can be captured using unobtrusive wearable accelerometers. Consistent with digital phenotyping approaches^71^, submovement-level analysis may help characterize subtle sensorimotor alterations through observable motor behavior, particularly among subclinical movement features that are otherwise difficult to characterize. Although these measures are not clinical endpoints per se^73,74^, they may contribute to early diagnosis, risk stratification, and treatment monitoring, all of that originating from a precise characterization of sensorimotor disfunction in psychiatry.

### 4.1 Limitations and future directions

Limitations should be considered when interpreting these findings. Group-level differences in submovement dynamics were robust, but the sample size was relatively small, and needs replication, possibly extending the methodology to individuals at clinical high-risk for psychosis, as well as antipsychotic-naïve patients. Longitudinal designs will be important to ascertain whether alterations in submovement structure evolve in parallel to disorder progression. Future studies should examine the relationship between submovement abnormalities and Neurological Soft Signs, especially regarding differential sensitivity detecting sensorimotor abnormalities. Finally, future work should examine the relationship between submovement abnormalities and broader interpersonal and social functioning.

## Ethical considerations

The study was approved by the Ethics Committee of *“Comitato Etico di Area Vasta Emilia Centro”,* protocol number EM255-2020-UniFe/170592-EM. All procedures were carried out in accordance with the ethical standards of the institutional research committee and with the 1964 Declaration of Helsinki. All participants provided written informed consent after receiving a verbal and written explanation of the task, procedures, and were informed of their right to withdraw at any time without consequences. Capacity to provide consent was assessed by the treating psychiatrist prior to enrolment, and only patients judged clinically stable and able to understand the study information were approached. Furthermore, recruitment and consent were overseen by clinical staff of the Psychiatric Unit of Sant’Anna Hospital, and all data were pseudonymized at collection. No follow-up data were collected, as the study used a single-session data collection. No animal subjects were involved.

## Acknowledgements

We would like to thank the staff of the Psychiatric Unit of Sant’Anna Hospital in Ferrara for welcoming us and supporting during data collection. We would also like to thank Matilde Pardi for her valuable contribution as a master’s student.

## Contributors

FP: investigation, formal analysis, writing-original draft, writing-review and editing. AT: conceptualization, supervision, writing-review and editing. AM: investigation, clinical recruitment, data collection, writing-review and editing. CG: investigation, writing-review and editing. GN: writing-review and editing. GADB: clinical recruitment, data collection, writing-review and editing. FT: writing-review and editing. GMG: writing-review and editing. MGN: writing-review and editing. LG: writing-review and editing. LF: writing-review and editing. MBM: conceptualization, clinical recruitment, supervision, writing-review and editing. AD: funding acquisition, conceptualization, supervision, writing-review and editing.

All authors read and approved the final version of the manuscript and agree to be accountable for all aspects of the work in ensuring that questions related to the accuracy or integrity of any part of the work are appropriately investigated and resolved. All persons designated as authors qualify for authorship, and all those who qualify for authorship are listed.

## Availability of data and materials

Data from this study will be made available on a public repository upon acceptance for publication.

