## Supplementary Material for "Microscopic Motor Alterations in Psychosis and Chronic Cannabis Use"

### Supplementary materials

#### 1. Demographic and clinical data

| Participants’ specific demographic data | | | |
|---|---|---|---|
| | **Group** | **Sex** | **Age** |
| S19 | HC | F | 44 |
| S20 | HC | F | 44 |
| S21 | HC | F | 21 |
| S22 | HC | F | 51 |
| S23 | HC | F | 23 |
| S24 | HC | F | 24 |
| S27 | HC | F | 21 |
| S28 | HC | M | 32 |
| S29 | HC | M | 27 |
| S30 | HC | F | 25 |
| S31 | HC | F | 43 |
| S33 | HC | F | 47 |
| S34 | HC | F | 23 |
| S40 | HC | M | 31 |
| S53 | HC | M | 29 |
| S54 | HC | M | 28 |
| S55 | HC | M | 23 |
| S56 | HC | M | 34 |
| S06 | HC-CU | F | 25 |
| S07 | HC-CU | M | 40 |
| S08 | HC-CU | M | 26 |
| S09 | HC-CU | M | 23 |
| S10 | HC-CU | M | 24 |
| S18 | HC-CU | M | 26 |
| S35* | HC-CU | M | 20 |
| S36 | HC-CU | F | 22 |
| S37 | HC-CU | F | 25 |
| S38 | HC-CU | F | 26 |
| S39 | HC-CU | M | 25 |
| S41 | HC-CU | M | 25 |
| S42 | HC-CU | M | 29 |
| S43 | HC-CU | F | 20 |
| S45 | HC-CU | F | 23 |
| S48 | HC-CU | M | 26 |

| | | | |
|---|---|---|---|
| S49 | HC-CU | M | 23 |
| S51 | HC-CU | M | 24 |
| S52 | HC-CU | M | 24 |
| S57 | HC-CU | M | 23 |
| S58 | HC-CU | M | 21 |
| S01 | PwP | M | 51 |
| S02 | PwP | F | 51 |
| S03 | PwP | F | 25 |
| S04 | PwP | M | 54 |
| S05 | PwP | F | 52 |
| S11 | PwP | F | 18 |
| S12 | PwP | M | 36 |
| S13 | PwP | F | 53 |
| S14 | PwP | F | 18 |
| S15 | PwP | F | 33 |
| S16 | PwP | F | 32 |
| S17 | PwP | F | 20 |
| S25 | PwP | F | 20 |
| S26 | PwP | M | 26 |
| S32 | PwP | M | 25 |
| S44 | PwP | M | 28 |
| S46 | PwP | M | 36 |
| S47* | PwP | M | 47 |
| S50* | PwP | M | 19 |

**Table S.1.** Participants' specific demographics. * S35, S47, and S50 were included in the study but kinematic data were unusable.

| ID | Regular use of cannabis | Chlorpromazine-equivalent mg |
|---|---|---|
| S01 | 1 | 4319 |
| S02 | 0 | 1092 |
| S03 | 1 | 5894 |
| S04 | 1 | 4158 |
| S05 | 0 | 500 |
| S11 | 1 | 40 |
| S12 | 1 | 1880 |
| S13 | 0 | 2760 |
| S14 | 0 | 385 |
| S15 | 0 | 71 |
| S16 | 0 | 20 |
| S17 | 0 | 159 |
| S25 | 0 | 1063 |
| S26 | 1 | 656 |
| S32 | 1 | 225 |
| S44 | 1 | 620 |
| S46 | 0 | 6577 |
| S47* | 0 | 1872 |
| S50* | 0 | 622 |

**Table S.2.** PwP group participants' specific data concerning chlorpromazine-equivalent dosage (mg) and regular use (yes = 1; no = 0) of cannabis. *S47, and S50 were included in the study but kinematic data were unusable.

#### 2. Time-locked kinematics and submovement probability change in the Biological anti-phase condition

The biological anti-phase condition showed a reduced submovement modulation and no structured post-submovement probability component (Figure S.1). Furthermore, the comparison between time-locked and surrogate probability distribution in this condition, revealed no significant differences at any timepoint.

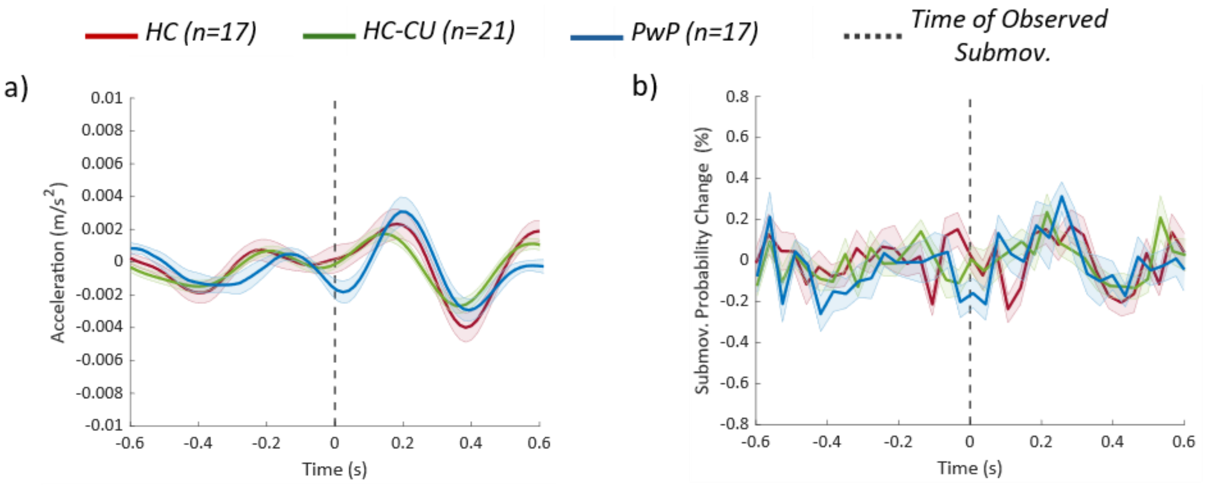


**Figure S.1.** Submovement-locked kinematics and probability across groups in the biological anti-phase condition. Acceleration profiles (a) and, probability of submovement occurrence (b), expressed as percentage deviation from the mean over the ±0.6 s interval.

#### 3. Point-by-point comparison between real and surrogate data

| Time (s) | FDR-corrected p-values | Raw p-values |
|---|---|---|
| -.212 | .010 | .003 |
| -.176 | .002 | 2.351e-04 |
| -.141 | .024 | .008 |
| -.035 | .037 | .015 |
| 0 | .010 | .003 |
| .035 | .002 | 4.951e-04 |
| .141 | .019 | .006 |
| .176 | 2.752e-04 | 1.573e-05 |
| .212 | .002 | 4.023e-04 |
| .318 | 3.824e-06 | 1.093e-07 |
| .353 | .002 | 2.932e-04 |
| .388 | .003 | 7.113e-04 |
| .494 | .045 | .019 |
| .529 | .026 | .010 |
| .6 | .002 | 3.563e-04 |

**Table S.3.** Real (time-locked) vs surrogate (shuffled time-locked) significant time Points and adjusted p-values in the HC Group during the in-phase condition. Note: only timepoints whose p-values are survived to the FDR correction method have been reported, along with their original raw p-value.

| Time (s) | FDR-corrected p-values | Raw p-values |
|---|---|---|
| -.600 | .006 | .001 |
| -.388 | .033 | .016 |
| -.212 | .001 | 1.591e-04 |
| -.141 | .002 | 2.796e-04 |
| .000 | .019 | .008 |
| .035 | .005 | 9.347e-04 |
| .106 | .033 | .015 |
| .141 | .002 | 2.428e-04 |
| .176 | 6.564e-05 | 3.751e-06 |
| .212 | .015 | .005 |
| .282 | .019 | .008 |
| .318 | 7.195e-05 | 6.167e-06 |
| .353 | .007 | .002 |
| .388 | 6.564e-05 | 2.284e-06 |
| .424 | .033 | .016 |
| .494 | .009 | .002 |
| .529 | .019 | .007 |

**Table S.4.** Real (time-locked) vs surrogate (shuffled time-locked) significant time Points and adjusted p-values in the HC-CU Group during the in-phase condition. Note: only timepoints whose p-values are survived to the FDR correction method have been reported, along with their original raw p-value.

| Time (s) | FDR-corrected p-values | Raw p-values |
|---|---|---|
| -.212 | .043 | .007 |
| -.176 | .043 | .011 |
| .000 | .043 | .010 |
| .141 | .026 | .002 |
| .176 | .040 | .005 |
| .212 | .043 | .010 |
| .318 | .043 | .012 |
| .388 | .026 | .001 |
| .424 | .043 | .010 |
| .529 | .020 | 5.742e-04 |

**Table S.5.** Real (time-locked) vs surrogate (shuffled time-locked) significant time Points and adjusted p-values in the PwP Group during the in-phase condition. Note: only timepoints whose p-values are survived to the FDR correction method have been reported, along with their original raw p-value.

Following the point-by-point *t*-tests comparing real (time-locked) and surrogate (shuffled time-locked) data none of the groups (HC, HC-CU, PwP) showed timepoints surviving the FDR correction in the Biological anti-phase condition. Although some uncorrected *p*-values approached significance, these effects did not remain after controlling for multiple comparisons.

#### 4. Relating submovement features with clinical and pharmacological variables

| Influence of cannabis on clinical variables | | | | |
|---|---|---|---|---|
| *Variable* | *Beta* | *p* | *R2* | *p_FDR* |
| PANSS-pos | 6.778 | .190 | .112 | .820 |
| PANSS-neg | -.708 | .854 | .002 | .854 |
| PANSS-glob | 1.139 | .824 | .003 | .854 |
| PANSS-tot | 7.208 | .547 | .025 | .820 |
| BPRS | 5.750 | .450 | .041 | .820 |
| Chlorpromazine | 821.000 | .451 | .038 | .820 |

**Table S.6.** Linear regression exploring the influence of cannabis on clinical variables.

| Influence of cannabis on submovement characteristics | | | | |
|---|---|---|---|---|
| *Variable* | *Beta* | *p* | *R2* | *p_FDR* |
| Mean-I-SM-I-Sinusoidal-in-phase | .006 | .100 | .170 | .251 |
| Mean-I-SM-I-Biological-in-phase | .002 | .723 | .009 | .876 |
| Mean-I-SM-I-Sinusoidal-anti-phase | -.001 | .774 | .006 | .876 |
| SD-I-SM-I-Sinusoidal-in-phase | -.013 | .009 | .408 | .078 |
| SD-I-SM-I-Biological-in-phase | -.012 | .083 | .187 | .251 |
| SD-I-SM-I-Sinusoidal-anti-phase | -.007 | .069 | .204 | .251 |
| Acceleration-Peak-Amplitude | .001 | .280 | .077 | .467 |
| Acceleration-Peak-Latency | .072 | .220 | .098 | .440 |
| Probability-Change-Peak-Amplitude | .010 | .907 | .001 | .907 |
| Probability-Change-Peak-Latency | -.016 | .788 | .005 | .876 |

**Table S.7.** Linear regression exploring the influence of cannabis on submovement characteristics.

| Zero-order correlations (no influence of cannabis) | | | | |
|---|---|---|---|---|
| *Predictor* | *MovementVariable* | *r0* | *p0* | *p0_FDR* |
| PANSS-pos | Mean-I-SM-I-Sinusoidal-in-phase | .499 | .042 | .311 |
| PANSS-pos | Mean-I-SM-I-Biological-in-phase | .604 | .010 | .310 |
| PANSS-pos | Mean-I-SM-I-Sinusoidal-anti-phase | .310 | .226 | .664 |
| PANSS-pos | SD-I-SM-I-Sinusoidal-in-phase | -.008 | .977 | .993 |
| PANSS-pos | SD-I-SM-I-Biological-in-phase | .294 | .252 | .664 |
| PANSS-pos | SD-I-SM-I-Sinusoidal-anti-phase | .128 | .624 | .814 |
| PANSS-pos | Acceleration-Peak-Amplitude | .160 | .539 | .770 |
| PANSS-pos | Acceleration-Peak-Latency | .181 | .486 | .749 |
| PANSS-pos | Probability-Change-Peak-Amplitude | .337 | .186 | .657 |
| PANSS-pos | Probability-Change-Peak-Latency | .001 | .998 | .998 |
| PANSS-neg | Mean-I-SM-I-Sinusoidal-in-phase | .246 | .342 | .664 |
| PANSS-neg | Mean-I-SM-I-Biological-in-phase | .134 | .607 | .814 |
| PANSS-neg | Mean-I-SM-I-Sinusoidal-anti-phase | .175 | .502 | .749 |
| PANSS-neg | SD-I-SM-I-Sinusoidal-in-phase | .034 | .896 | .959 |
| PANSS-neg | SD-I-SM-I-Biological-in-phase | .062 | .812 | .921 |
| PANSS-neg | SD-I-SM-I-Sinusoidal-anti-phase | .257 | .320 | .664 |
| PANSS-neg | Acceleration-Peak-Amplitude | .240 | .354 | .664 |
| PANSS-neg | Acceleration-Peak-Latency | -.271 | .292 | .664 |
| PANSS-neg | Probability-Change-Peak-Amplitude | .458 | .064 | .430 |
| PANSS-neg | Probability-Change-Peak-Latency | .257 | .319 | .664 |
| PANSS-glob | Mean-I-SM-I-Sinusoidal-in-phase | .572 | .016 | .310 |
| PANSS-glob | Mean-I-SM-I-Biological-in-phase | .509 | .037 | .311 |
| PANSS-glob | Mean-I-SM-I-Sinusoidal-anti-phase | .296 | .249 | .664 |
| PANSS-glob | SD-I-SM-I-Sinusoidal-in-phase | .214 | .410 | .676 |
| PANSS-glob | SD-I-SM-I-Biological-in-phase | .329 | .197 | .657 |
| PANSS-glob | SD-I-SM-I-Sinusoidal-anti-phase | .229 | .376 | .664 |
| PANSS-glob | Acceleration-Peak-Amplitude | .056 | .832 | .921 |
| PANSS-glob | Acceleration-Peak-Latency | .233 | .369 | .664 |
| PANSS-glob | Probability-Change-Peak-Amplitude | .331 | .194 | .657 |
| PANSS-glob | Probability-Change-Peak-Latency | .024 | .928 | .960 |
| PANSS-tot | Mean-I-SM-I-Sinusoidal-in-phase | .544 | .024 | .310 |
| PANSS-tot | Mean-I-SM-I-Biological-in-phase | .527 | .030 | .310 |
| PANSS-tot | Mean-I-SM-I-Sinusoidal-anti-phase | .319 | .211 | .664 |
| PANSS-tot | SD-I-SM-I-Sinusoidal-in-phase | .099 | .706 | .851 |
| PANSS-tot | SD-I-SM-I-Biological-in-phase | .290 | .258 | .664 |
| PANSS-tot | SD-I-SM-I-Sinusoidal-anti-phase | .237 | .360 | .664 |
| PANSS-tot | Acceleration-Peak-Amplitude | .171 | .511 | .749 |
| PANSS-tot | Acceleration-Peak-Latency | .093 | .724 | .851 |
| PANSS-tot | Probability-Change-Peak-Amplitude | .437 | .079 | .475 |
| PANSS-tot | Probability-Change-Peak-Latency | .093 | .723 | .851 |
| BPRS | Mean-I-SM-I-Sinusoidal-in-phase | .547 | .028 | .310 |
| BPRS | Mean-I-SM-I-Biological-in-phase | .540 | .031 | .310 |
| BPRS | Mean-I-SM-I-Sinusoidal-anti-phase | .379 | .148 | .657 |
| BPRS | SD-I-SM-I-Sinusoidal-in-phase | .027 | .921 | .960 |
| BPRS | SD-I-SM-I-Biological-in-phase | .258 | .334 | .664 |
| BPRS | SD-I-SM-I-Sinusoidal-anti-phase | .272 | .308 | .664 |
| BPRS | Acceleration-Peak-Amplitude | .218 | .417 | .676 |
| BPRS | Acceleration-Peak-Latency | .063 | .817 | .921 |

| | | | | |
|---|---|---|---|---|
| BPRS | Probability-Change-Peak-Amplitude | .349 | .185 | .657 |
| BPRS | Probability-Change-Peak-Latency | .144 | .596 | .814 |
| Chlorpromazine | Mean-I-SM-I-Sinusoidal-in-phase | -.052 | .844 | .921 |
| Chlorpromazine | Mean-I-SM-I-Biological-in-phase | .331 | .195 | .657 |
| Chlorpromazine | Mean-I-SM-I-Sinusoidal-anti-phase | .263 | .307 | .664 |
| Chlorpromazine | SD-I-SM-I-Sinusoidal-in-phase | -.411 | .101 | .553 |
| Chlorpromazine | SD-I-SM-I-Biological-in-phase | .129 | .621 | .814 |
| Chlorpromazine | SD-I-SM-I-Sinusoidal-anti-phase | .117 | .654 | .835 |
| Chlorpromazine | Acceleration-Peak-Amplitude | .337 | .186 | .657 |
| Chlorpromazine | Acceleration-Peak-Latency | -.221 | .393 | .674 |
| Chlorpromazine | Probability-Change-Peak-Amplitude | .171 | .512 | .749 |
| Chlorpromazine | Probability-Change-Peak-Latency | -.110 | .673 | .842 |

**Table S.8.** Zero-order correlation exploring the relationship between submovement features and clinical or pharmacological variables unadjusted for cannabis use.

| Partial_correlations_controlling_for_cannabis_use | | | | |
|---|---|---|---|---|
| *Predictor* | *MovementVariable* | *r_partial* | *p_partial* | *p_partial_ FDR* |
| PANSS-pos | Mean-I-SM-I-Sinusoidal-in-phase | .421 | .105 | .519 |
| PANSS-pos | Mean-I-SM-I-Biological-in-phase | .610 | .012 | .356 |
| PANSS-pos | Mean-I-SM-I-Sinusoidal-anti-phase | .357 | .175 | .519 |
| PANSS-pos | SD-I-SM-I-Sinusoidal-in-phase | .284 | .286 | .520 |
| PANSS-pos | SD-I-SM-I-Biological-in-phase | .516 | .041 | .356 |
| PANSS-pos | SD-I-SM-I-Sinusoidal-anti-phase | .332 | .209 | .519 |
| PANSS-pos | Acceleration-Peak-Amplitude | .074 | .784 | .888 |
| PANSS-pos | Acceleration-Peak-Latency | .085 | .753 | .869 |
| PANSS-pos | Probability-Change-Peak-Amplitude | .347 | .188 | .519 |
| PANSS-pos | Probability-Change-Peak-Latency | .026 | .925 | .957 |
| PANSS-neg | Mean-I-SM-I-Sinusoidal-in-phase | .292 | .273 | .520 |
| PANSS-neg | Mean-I-SM-I-Biological-in-phase | .140 | .606 | .757 |
| PANSS-neg | Mean-I-SM-I-Sinusoidal-anti-phase | .172 | .525 | .727 |
| PANSS-neg | SD-I-SM-I-Sinusoidal-in-phase | .005 | .986 | .995 |
| PANSS-neg | SD-I-SM-I-Biological-in-phase | .046 | .866 | .943 |
| PANSS-neg | SD-I-SM-I-Sinusoidal-anti-phase | .264 | .324 | .529 |
| PANSS-neg | Acceleration-Peak-Amplitude | .264 | .323 | .529 |
| PANSS-neg | Acceleration-Peak-Latency | -.270 | .312 | .529 |
| PANSS-neg | Probability-Change-Peak-Amplitude | .460 | .073 | .486 |
| PANSS-neg | Probability-Change-Peak-Latency | .254 | .342 | .539 |
| PANSS-glob | Mean-I-SM-I-Sinusoidal-in-phase | .602 | .014 | .356 |
| PANSS-glob | Mean-I-SM-I-Biological-in-phase | .506 | .045 | .356 |
| PANSS-glob | Mean-I-SM-I-Sinusoidal-anti-phase | .301 | .257 | .520 |
| PANSS-glob | SD-I-SM-I-Sinusoidal-in-phase | .327 | .216 | .519 |
| PANSS-glob | SD-I-SM-I-Biological-in-phase | .393 | .132 | .519 |
| PANSS-glob | SD-I-SM-I-Sinusoidal-anti-phase | .287 | .281 | .520 |
| PANSS-glob | Acceleration-Peak-Amplitude | .041 | .880 | .943 |
| PANSS-glob | Acceleration-Peak-Latency | .226 | .399 | .584 |
| PANSS-glob | Probability-Change-Peak-Amplitude | .330 | .212 | .519 |
| PANSS-glob | Probability-Change-Peak-Latency | .028 | .918 | .957 |
| PANSS-tot | Mean-I-SM-I-Sinusoidal-in-phase | .532 | .034 | .356 |
| PANSS-tot | Mean-I-SM-I-Biological-in-phase | .521 | .038 | .356 |
| PANSS-tot | Mean-I-SM-I-Sinusoidal-anti-phase | .336 | .203 | .519 |
| PANSS-tot | SD-I-SM-I-Sinusoidal-in-phase | .263 | .326 | .529 |
| PANSS-tot | SD-I-SM-I-Biological-in-phase | .402 | .122 | .519 |
| PANSS-tot | SD-I-SM-I-Sinusoidal-anti-phase | .349 | .185 | .519 |
| PANSS-tot | Acceleration-Peak-Amplitude | .135 | .619 | .758 |
| PANSS-tot | Acceleration-Peak-Latency | .046 | .865 | .943 |
| PANSS-tot | Probability-Change-Peak-Amplitude | .438 | .090 | .519 |
| PANSS-tot | Probability-Change-Peak-Latency | .105 | .698 | .837 |
| BPRS | Mean-I-SM-I-Sinusoidal-in-phase | .519 | .047 | .356 |
| BPRS | Mean-I-SM-I-Biological-in-phase | .526 | .044 | .356 |
| BPRS | Mean-I-SM-I-Sinusoidal-anti-phase | .403 | .136 | .519 |
| BPRS | SD-I-SM-I-Sinusoidal-in-phase | .207 | .458 | .655 |
| BPRS | SD-I-SM-I-Biological-in-phase | .372 | .172 | .519 |
| BPRS | SD-I-SM-I-Sinusoidal-anti-phase | .400 | .139 | .519 |
| BPRS | Acceleration-Peak-Amplitude | .161 | .565 | .745 |

| | | | | |
|---|---|---|---|---|
| BPRS | Acceleration-Peak-Latency | -.002 | .995 | .995 |
| BPRS | Probability-Change-Peak-Amplitude | .346 | .206 | .519 |
| BPRS | Probability-Change-Peak-Latency | .159 | .571 | .745 |
| Chlorpromazine | Mean-I-SM-I-Sinusoidal-in-phase | -.148 | .584 | .745 |
| Chlorpromazine | Mean-I-SM-I-Biological-in-phase | .320 | .227 | .520 |
| Chlorpromazine | Mean-I-SM-I-Sinusoidal-anti-phase | .284 | .286 | .520 |
| Chlorpromazine | SD-I-SM-I-Sinusoidal-in-phase | -.378 | .148 | .519 |
| Chlorpromazine | SD-I-SM-I-Biological-in-phase | .242 | .367 | .565 |
| Chlorpromazine | SD-I-SM-I-Sinusoidal-anti-phase | .235 | .380 | .570 |
| Chlorpromazine | Acceleration-Peak-Amplitude | .300 | .259 | .520 |
| Chlorpromazine | Acceleration-Peak-Latency | -.304 | .253 | .520 |
| Chlorpromazine | Probability-Change-Peak-Amplitude | .168 | .533 | .727 |
| Chlorpromazine | Probability-Change-Peak-Latency | -.099 | .716 | .842 |

**Table S.9.** Partial correlations adjusting for cannabis use, exploring the relationship between submovement features and clinical or pharmacological variables adjusted for cannabis use.

#### 5. Control analysis of task performance

To ensure that participants across all groups performed the task with comparable levels of basic synchronicity, the mean absolute difference between participants' and HKT's timestamps of movement onset/offset were analysed using a mixed-design ANOVA, with Group (HC, HC-CU, and PwP) as the between-subject factor and Condition (biological in-phase, biological anti-phase) as the within-subject factor, implemented in JASP (Version 0.19.3, https://jasp-stats.org/). The analysis revealed no significant main effect of Group ($F(2,52) = .541$, $p= .585$, $\eta^2 = .014$), indicating that the general level of synchronicity did not differ between HC, HC-CU, and PwP. Similarly, the main effect of Condition was not significant ($F(1,52) = 1.868$, $p = .178$, $\eta^2 = .009$). Finally, the Group × Condition interaction did not reach significance ($F(2,52) = 2.369$, p = .104, $\eta^2 = .023$). These results confirm that all groups were able to follow the start and stop of the stimulus with similar accuracy across conditions.
